# Oscillatory Networks of High-Level Mental Alignment: A Perspective-Taking MEG Study

**DOI:** 10.1101/198747

**Authors:** R.A Seymour,, H. Wang,, G. Rippon,, K. Kessler,

## Abstract

Mentally imagining another’s perspective is a high-level social process, reliant on manipulating internal representations of the self in an embodied manner. Recently Wang et al., (1) showed that theta-band (3-7Hz) brain oscillations within the right temporo-parietal junction (rTPJ) and brain regions coding for motor/body schema contribute to the process of perspective-taking. Using a task requiring participants to engage in embodied perspective-taking, we set out to unravel the extended functional brain network and its connections in detail. We found that increasing the angle of disparity between self and other perspective was accompanied by longer reaction times and increases in theta power within rTPJ, right lateral pre-frontal cortex (PFC) and right anterior cingulate cortex (ACC). Using nonparametric Granger-causality, we showed that during later stages of perspective-taking, the lateral PFC and ACC exert top-down influences over rTPJ, indicative of executive control processes required for managing conflicts between self and other perspectives. Finally, we quantified patterns of whole-brain phase coupling (imaginary coherence) in relation to rTPJ during high-level perspective taking. Results suggest that rTPJ increases its theta-band phase synchrony with brain regions involved in mentalizing and regions coding for motor/body schema; whilst decreasing its synchrony to visual regions. Implications for neurocognitive models are discussed, and it is proposed that rTPJ acts as a ‘hub’ to route bottom-up visual information to internal representations of the self during perspective-taking, co-ordinated by theta-band oscillations. The self is then projected onto the other’s perspective via embodied motor/body schema transformations, regulated by top-down cingulo-frontal activity.

**Significance Statement:** High-level social processing, such as the ability to imagine another’s visuospatial experience of the world (perspective taking), is a core part of what makes us human. Building on a substantial body of converging previous evidence, our study reveals how concerted activity across the cortex in low frequencies (theta: 3-7 Hz) implements this crucial human process. We found that oscillatory power and connectivity (imaginary coherence, nonparametric Granger causality) at theta frequency linked functional sub-networks of executive control, mentalizing, and sensorimotor/body schema via a main hub located in the right temporo-parietal junction (rTPJ). Our findings inform neurocognitive models of social cognition by describing the co-ordinated changes in brain network connectivity, mediated by theta oscillations, during perspective-taking.

## Introduction

Humans possess highly developed social skills that allow us to imagine what others might be experiencing, thinking or feeling to an extent not shared by other species (2). The question of what separates us from other species has been the subject of substantial research in comparative psychology and cognitive neuroscience, and while significant headway has been made with respect to what skills make us special (3–5) and which parts of our brain have evolved to cope with sophisticated “mentalizing”, i.e., reading of others’ minds (6, 7), much less is known about the actual brain network dynamics that implement these social skills.Here we set out to investigate the large-scale, distributed but synchronised neural activity that gives rise to a person’s understanding of another’s visuospatial experience of the world: a process termed perspective taking (P-taking).

Mentally imagining another’s perspective is a high-level social process, but recent behavioural experiments suggest that P-taking is still grounded in the cortical posture and action representations of the observer. Using posture manipulations, several studies (1, 8–11) have shown that P-taking engages large parts of the neuronal bases of the body schema, i.e.the cortical correlates of the internal representation of the body (12, 13), in the form of a simulated rotation of the embodied self into another’s orientation and perspective (1, 9, 10). In other words, humans literally “put themselves” into another’s viewpoint to understand their perspective.

Note that such embodied P-*taking* must be distinguished from so-called perspective *tracking* (P-tracking). While both processes involve judgements about another’s perspective, P-*tracking*, in contrast to P-*taking*, merely requires an observer to understand what another can or cannot perceive (e.g. what is occluded and what is visible to them). The two forms of perspective processing have been related to different developmental stages (14–16) (P-tracking: ∼2 years; P-taking ∼ 4-5 years) and P-tracking, in contrast to P-taking, has been observed in other species such as apes and corvids (17, 18). Finally, while P-taking engages an embodied mental rotation of the self into another’s viewpoint, P-tracking seems to rely on inferring another’s line of sight, in other words, whether their line of sight towards a target is disrupted or not (1, 8, 19).

The neural correlates of embodied simulation during P-taking were recently investigated by Wang and colleagues (1) using Magnetoencephalography (MEG, Expt. 1) and converging effects were found in the right posterior temporo-parietal junction (pTPJ) for cognitive effort of P-taking (amount of angular disparity between self vs. other’s viewpoint) and for embodied processing (posture congruence) during P-taking (but not for P-tracking). The crucial role of right pTPJ for P-taking was further confirmed via transcranial magnetic stimulation (TMS) interference (1). Wang et al. further reported that low frequency theta oscillations (3-7 Hz) were the prominent neural code in pTPJ, whilst Gooding-Williams et al., (11) used repetitive TMS entrainment over pTPJ to show that TMS pulses administered at theta frequency (6Hz) accelerated P-taking, while alpha (10Hz) entrainment slowed P-taking down. TPJ-theta could therefore be the relevant neural frequency to enable phase-coupling within a wider mentalizing network.

These results are consistent with the neural correlates of P-taking reported using fMRI – two meta-analyses (20, 21) have suggested that the core areas of activation include bilateral TPJ and ventro-medial pre-frontal cortex (vmPFC). The posterior division of the TPJ (22, 23) in particular, has been reliably linked to P-taking and more generally to “mentalizing” (representing other’s mental states) (21, 24), as well as to so-called spontaneous “out-of-body experiences” (OBE) (25). During an OBE individuals experience the sensation that the self has moved to a different physical location than their body, and this sensation often entails a translation as well as a rotation of perspective, similar to a deliberate perspective transformation during P-taking (26). The involvement of TPJ in OBEs (25) is of importance, as it corroborates the proposed link between embodied processing and high-level social mentalizing in TPJ (1, 25–27).

Whilst the TPJ is clearly important for embodied processing and P-taking, the region is also implicated in a range of cognitive operations, including spatial attention, social cognition and self/other distinctions. It has been suggested that more generally, the region acts as a major hub for information integration (22, 28), especially during higher-level cognitive processes relying upon internal representations, such as P-taking (1, 11, 22, 28). Indeed, the TPJ has extensive functional connectivity to many networks of the brain, including the fronto-parietal control (29), default mode (30), and ventral attention networks (23). We therefore hypothesised that the TPJ contributes to the process of embodied transformation through changes in patterns of whole-brain functional connectivity, via theta-band synchrony, as would be predicted from the region’s role as a network hub (22, 28, 31). However, investigations of P-taking using connectivity analysis, e.g. in form of frequency-specific phase-coupling, are scarce. To our knowledge, only one study to date (32) has reported enhanced theta phase-coherence between right TPJ and ventromedial prefrontal cortex (vmPFC) in a condition that required participants to imagine another’s visual experience. The major aim of the current study was therefore to consolidate the crucial role of pTPJ theta oscillations in P-taking by means of advanced network analyses.

In addition to the TPJ, Wang et al., reported increases in theta-band power for the lateral PFC during the cognitive effort of P-taking (1). Activity within this region during social cognition has been argued to reflect high-level reasoning and working memory processes recruited more generally during complex P-taking and mentalizing tasks (21). However, there is emerging evidence that frontal activity in lateral PFC but also in the anterior cingulate cortex (ACC) could play a more nuanced role in P-taking by managing the conflict between self and other perspectives (32–34). This could potentially manifest as a direct connection between lateral PFC and the core mentalizing network (32) in TPJ and vmPFC (20, 21). We were therefore interested in whether the TPJ becomes functionally connected to various frontal and midline regions during P-taking (34), and crucially determining the direction of this connectivity.

In conclusion, we set out to consolidate previous findings regarding the crucial role of TPJ theta oscillations for generating the abstract social representations required for P-taking (1, 11, 32), while unravelling in detail the involved functional network in terms of dynamic oscillatory coupling between brain areas, using MEG. Based on the considerations above, we expected TPJ and (v)mPFC to form a mentalizing network synchronised via theta oscillations, related to generating the abstract representation of another’s perspective, while activation in parietal body-schema areas and sensorimotor cortex would reflect the required embodied transformation to generate this representation via rotation of the egocentric perspective (8–10). In addition, pACC and lPFC might play key roles in top-down executive control of the underlying embodied transformation and in managing the conflict between physical self and transformed self at the representational level.

## Results

Whilst undergoing MEG, participants (n=18) performed a P-taking task, adapted from Kessler & Rutherford (8), requiring participants to determine the location of a target dot in respect to an avatar’s visuospatial perspective. The target could either be to the left/right (LR) of the avatar (requiring P-taking) or visible/occluded (VO) from the avatar (requiring P-tracking), and presented either 160° or 60° from the participant’s perspective. This resulted in four experimental conditions: LR-160; LR-60; VO-160; VO-60 (see Fig. 1; Materials and Methods). This 2x2 design allowed us to disentangle P-taking from P-tracking and investigate the effect of an increased angle of disparity (we chose to use 160° and 60° based on the results of Wang et al., (1)), between self-perspective and other-perspective, which has been shown to lengthen reaction times during P-taking (8, 10).

**Figure 1:**
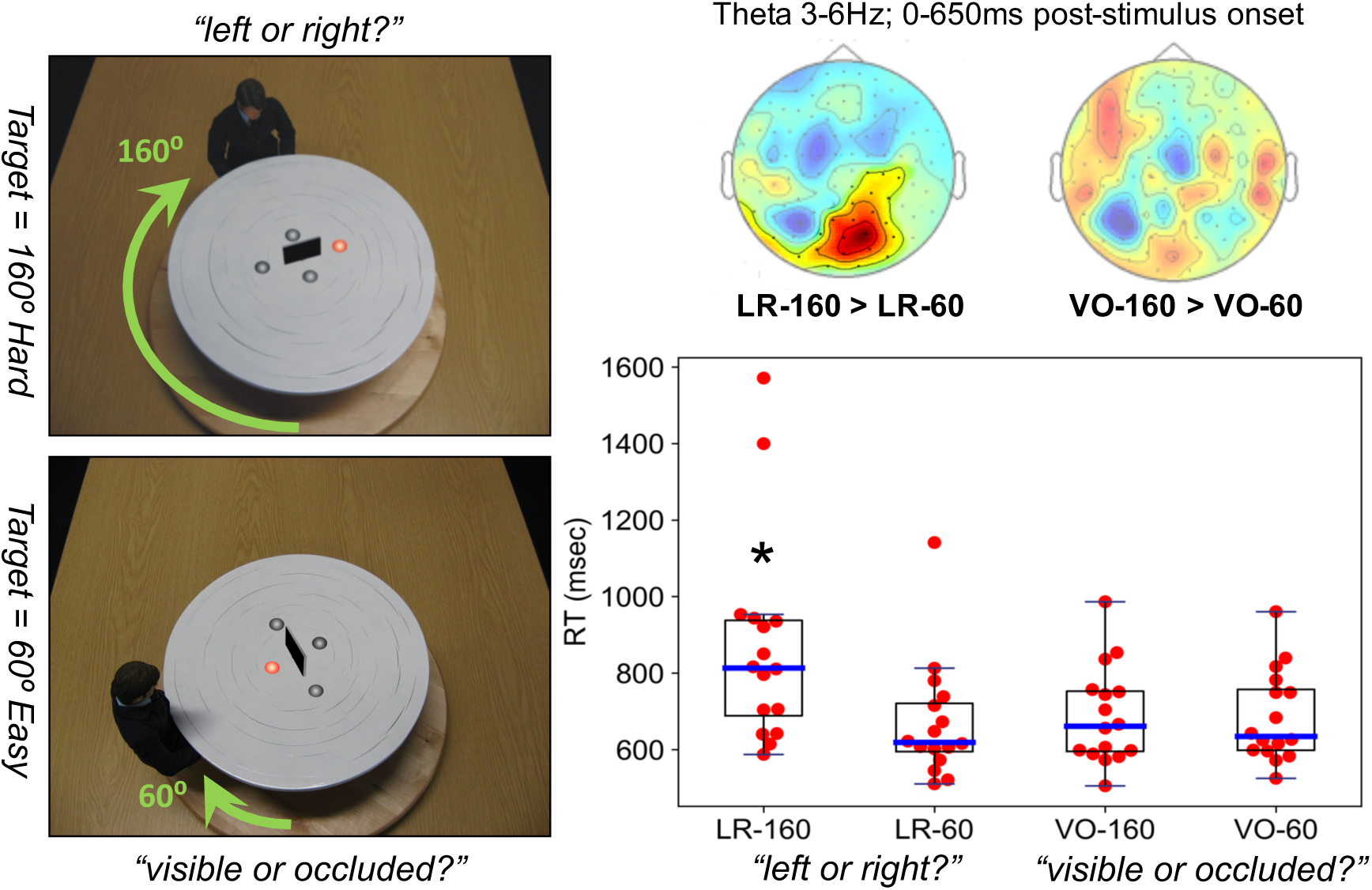
Left: Experimental paradigm (8) showing two example stimuli (see Materials and Methods for details). Bottom right: Boxplot of participants’ median reaction time (RT) in milliseconds for the two angular disparity conditions (160 vs. 60) of P-taking (L/R) and P-tracking (V/O), respectively. * = LR-160 is significantly different from all other conditions (p<.05). Top right: Sensor-level topoplots of theta activity (3-6Hz), showing a significant cluster (high visibility) for P-taking but not for P-tracking (further details in Materials and Methods and SI).

### Behavioural Results

For the four experimental conditions, median reaction times (RT) from each participant were entered into a one-way ANOVA (output detailed in Table S2). Results showed a main effect of experimental condition on RT, F(3,60) = 4.43, p=.007. Post-hoc tests revealed this was due to significantly longer RT for the LR-160 conditions compared with all other conditions (LR-60, p_tukey_= .013; VO-160, p_tukey_= .029; VO-60, p_tukey_= 026), replicating Kessler & Rutherford (8).

### Task-Related Changes in Theta Power

We used the open-source Fieldtrip toolbox and customised Matlab scripts for all MEG analyses (full details in Materials and Methods). Statistical analyses (at sensor and source level) were performed using cluster-based non-parametric permutation testing to correct for multiple comparisons across sensors/voxels (35). Using a data-driven approach from 1-30Hz, time-frequency results at the sensor-level (see Fig. 1 and Fig. S1) replicated the crucial role of theta oscillations in P-taking (1, 11). A significant positive cluster (p=0.03) was found at 3-6Hz, 0-650ms, when comparing angular disparities of 160° and 60° degrees for the L/R task. No significant clusters were found for the V/O task, i.e. P-tracking in the VO-160 vs VO-60 contrast.

To investigate the cortical sources underlying this effect of angular disparity, theta-band (3-6Hz) power was localised from 0-650ms post-stimulus onset separately for 160° and 60° trials, using the Dynamic Imaging of Coherent Sources (DICS) approach (see Materials and Methods). Baseline-corrected theta (3-6Hz) power was compared for LR-160 versus LR-60 trials and VO-160 versus VO-60 trials across a 0.8cm cortical grid (Fig. 2A; Supplementary Table S3). Results showed a significant (35) (p<.05) increase in theta power during LR-160 trials compared with LR-60 trials for right posterior temporo-parietal junction (pTPJ) spreading into the inter-parietal sulcus (IPS), for right lateral pre-frontal cortex (PFC) primarily overlapping with the inferior frontal gyrus (IFG) and for right anterior cingulate cortex (ACC). There was also a decrease in theta power in the LR-160 versus LR-60 condition in the left frontal pole (Table S3). For P-tracking (VO-task), there were increases in theta-power, compared with pre-trial baseline, within ventral occipital and temporo-parietal regions (Fig. S2). However, the VO-160 > VO-60 contrast showed no significant clusters (Fig. S3), indicating that angular disparity (160° versus 60°) only resulted in task-related increases in theta-band power for high-level P-taking and not for P-tracking.

**Figure 2:**
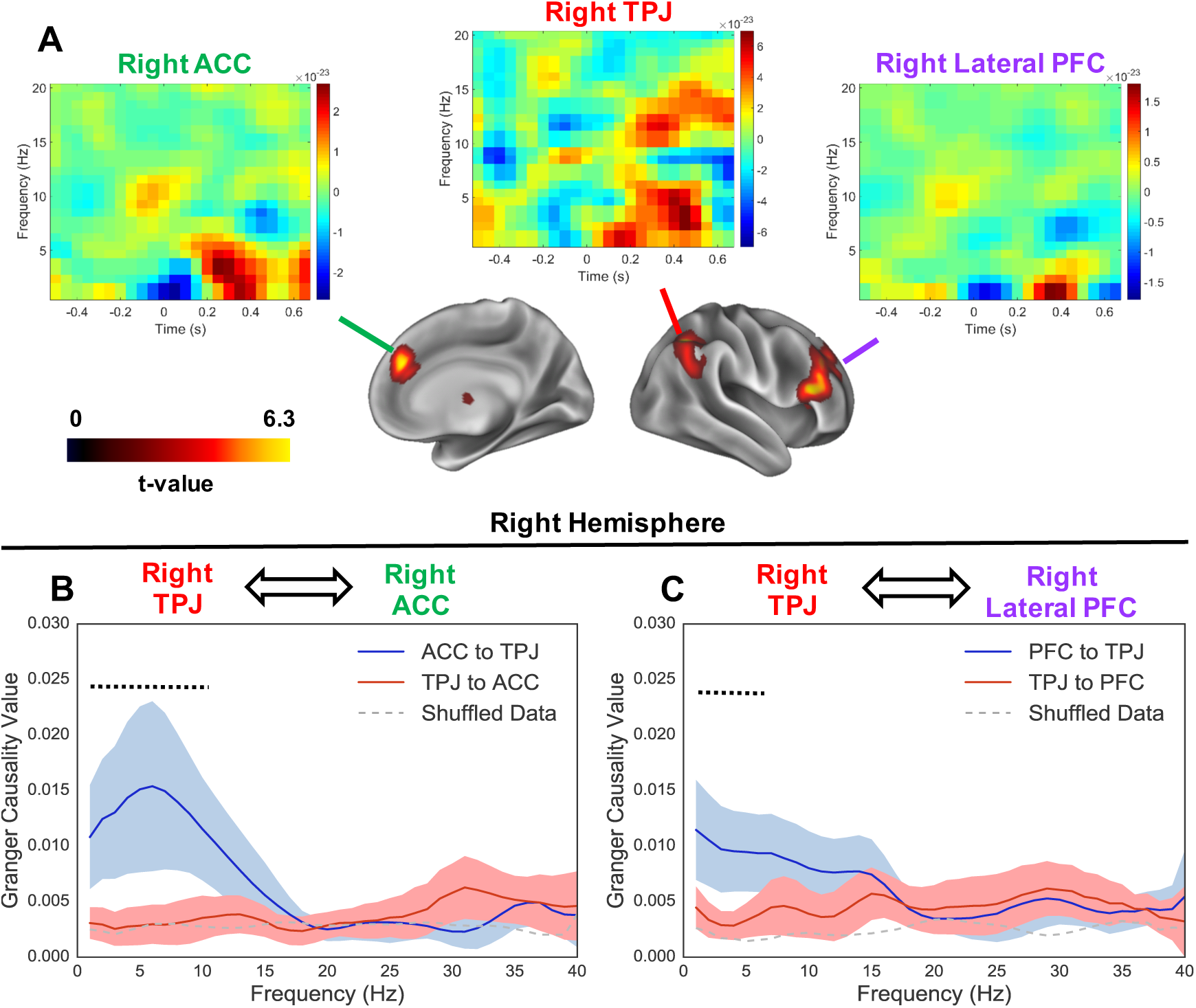
Theta power sources and directed connectivity. Panel A depicts brain plots showing statistical results (clusters with p<.05 are shown) of a whole-brain DICS theta power (3-6Hz) analysis for LR-160 > LR-60 contrast visualised using the Connectome Workbench software (36) (see Table S3 for a complete list of power sources). Plots at the top show time-frequency representations (LR-160 > LR-60 contrast) for three virtual electrodes (VE) placed in right ACC (MNI co-ordinates: [12, 36, 28]), right TPJ (MNI co-ordinates [40-58 36]) and right lateral PFC (MNI co-ordinates: [52,32,16]). Bottom plots show spectrally resolved non-parametric Granger causality (1-40Hz), computed between the right TPJ and (A) right ACC and (B) right lateral PFC, respectively. Results show an increase in Granger causality from both the right ACC (1-10Hz) and right PFC (1-5Hz) to the right TPJ. Shaded regions around each line represent 95% confidence intervals. The black dotted line above the plots represents Granger causality values passing a p<.05 threshold (see Methods for details). The grey dotted line in the plots shows shuffled data for comparison. Further explanations in the text.

### Virtual Electrode Time-Frequency Analysis

To further investigate the oscillatory signatures of high-level P-taking, time-courses for each trial were extracted from ‘virtual-electrodes’ in right TPJ, right ACC and right lateral PFC (see Materials & Methods for details). Low-frequency oscillatory power was then estimated between -0.65 to 0.65s post-stimulus using a Hanning taper, 0.05s sliding window. Results show very early and sustained theta power (3-6Hz) increases in the right TPJ (0-0.5s) for LR-160 versus LR-60 trials. Right lateral PFC delta/theta power (1-5Hz) and right ACC (1-5Hz) increases are more transient and begin from 0.2-0.5s post-stimulus onset. This suggests that the rTPJ is engaged throughout the process of embodied P-taking, whereas increases in theta power ACC and PFC occur later and more transiently.

### Granger Causality Analysis

To investigate directed functional connectivity during P-taking between the three main regions of interest (ROIs) identified in the source power analysis (rTPJ, rACC and rPFC), we employed non-parametric Granger causality (GC) on LR trials (0-0.65s post-stimulus onset) (37). GC is a statistical technique to estimate directionality between time-series, including human brain imaging data (37). GC values showed statistically significant (39) differences from fourier-scrambled time-series between two ROI pairs: rTPJ-rACC and rTPJ-rPFC. To investigate these effects further, we statistically compared GC values between each direction of the ROI pair (i.e. the granger causal influence to and from the rTPJ). Results showed an asymmetric increase in granger causal influence, directed top-down from right ACC between 1-10Hz, with a peak at 6Hz, (Fig. 2B, p=.009) and right PFC, between 1-6Hz, (Fig 2B, p=.04) to the right TPJ.

### Imaginary Coherence

To establish patterns of whole-brain functional connectivity accompanying right-hemisphere TPJ theta-band activity, we extracted source-level theta-band (5±2 Hz) phase relationships from the sensor-level cross-spectral density matrix using an adaptive spatial-filtering approach (38) (see Materials and Methods). A measure of phase synchrony between a right-TPJ seed (MNI co-ordinates: [40-58 36]) and every other voxel was calculated by projecting complex-valued coherency onto the imaginary axis, thus, reducing the influence of MEG field spread by removing instantaneous phase (39). The resulting coherency maps from the LR-160 and LR-60 conditions were first baseline-corrected, and then compared using cluster-based non-parametric permutation testing (35).

Results (Fig. 3) show a complex pattern of both increased and decreased theta-band phase synchrony during embodied P-taking. The main areas of decreased synchrony are located in the ventral occipitotemporal cortex (VOTC), overlapping with key regions of the ventral visual stream. There were also reductions in phase synchrony to the bilateral anterior temporal lobes (ATL). Increased phase synchrony was observed in bilateral medial PFC regions, posterior cingulate cortex (PCC), intra-parietal sulcus (IPS), supplementary motor area (SMA), posterior parietal cortex (PPC), and right supramarginal gyrus/sensorimotor cortex (SMC). These patterns of phase synchrony are unlikely to be driven by spurious connectivity from MEG field spread (40), as we opted to measure imaginary coherence (39), thereby removing effects in relation to instantaneous phase.

**Figure 3:**
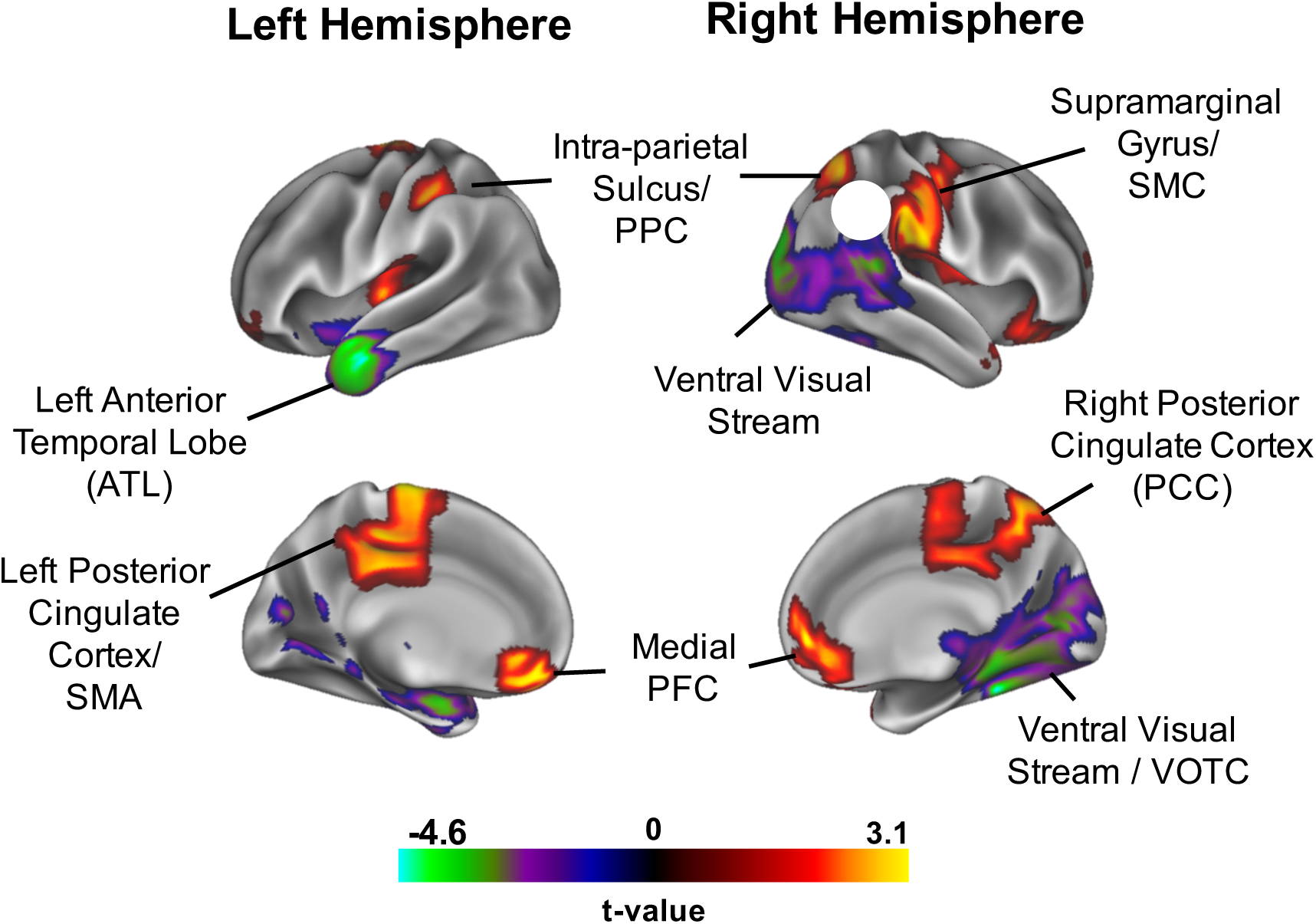
Results of a whole-brain imaginary coherence analysis in relation to a right TPJ seed (white circle) and for a LR-160 > LR-60 contrast, visualised using the Connectome Workbench software (36) (see Table S4 for a complete list of sources). Clusters of coherency increase/decrease passing a p<.05 threshold are shown (see Material and Methods). PPC = posterior parietal cortex; SMC = sensorimotor cortex; SMA = supplementary motor area; PFC = prefrontal cortex; VOTC = ventral occipitotemporal cortex.

## Discussion

This MEG study has investigated the oscillation-based functional connectivity between brain regions involved in our ability to take another person’s visuospatial perspective. Behavioural results replicated a substantial body of research showing significantly increased reaction time for higher angular disparity between the participant and avatar (160° versus 60°) for perspective-*taking* (P-taking) but not perspective-*tracking* (P-tracking) (1, 8–10). Greater angular disparity for P-taking was accompanied by increased theta power (3-6Hz) within the right TPJ/IPS and lateral PFC, replicating Wang et al., (1), as well as within the right ACC.

Importantly, this increase in theta-power for angular disparity was specific to P-taking and not P-tracking (Figs. 1, S1, S3). We therefore focused on network connectivity during P-taking and showed (Fig. 2) that there was an increase in Granger causal influence (37) from lateral PFC and right ACC *to* right TPJ, but not vice-versa, mediated by low frequency brain rhythms (1-10Hz). Finally, we examined how whole-brain patterns of theta-band (5±2 Hz) phase synchrony, quantified using imaginary coherence (39), varied in relation to right TPJ activity. Results (Fig. 3) suggest that with increasing angular disparity (160° versus 60°), the right TPJ increases its phase coupling to regions involved in theory of mind (41) (medial PFC, PCC) and body schema (12, 13) (SMC, PPC, SMA), but decreases its phase coupling to visual regions (VOTC) and to bilateral anterior temporal lobe (ATL). Overall, these results suggest a crucial role for TPJ as a hub that functionally connects mentalizing, executive, and body-representational networks via theta-band (3-7Hz) oscillations during high-level perspective taking (Fig. 4).

**Figure 4:**
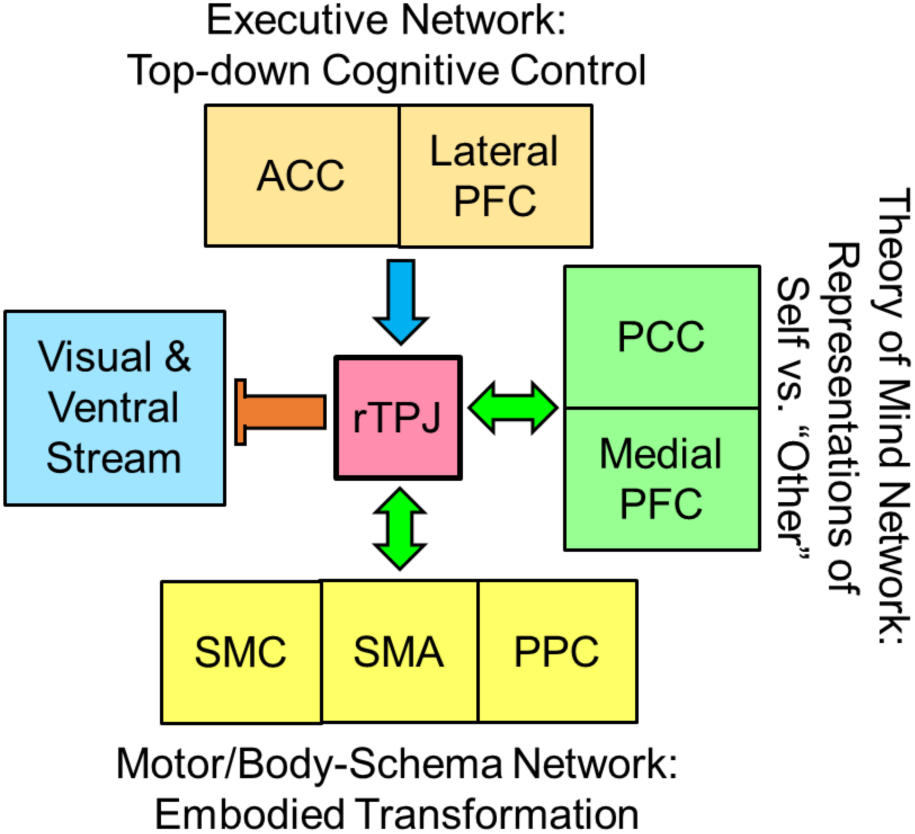
Proposed network underlying high-level perspective-taking (8, 10), linked by power and phase in the theta-band (3-7Hz). During initiation of embodied perspective-taking behaviour, early rTPJ activity co-ordinates connectivity decreases with visual regions, whilst increasing connectivity with regions involved in Theory of Mind, and Motor/Body-Schema. Increases in theta power within the lateral PFC and ACC reflect the top-down domain-general cognitive control processes for detecting and resolving the conflict between self and other perspectives (44).

### The role of the right TPJ in high-level perspective-taking

Results from this study suggest that the right TPJ (rTPJ) becomes increasingly engaged with the need for embodied mental alignment during P-taking i.e. as the disparity grows between our own and other’s perspectives (1, 11). Activity within the right TPJ is consistent with its role in establishing a sense of self (25), and crucially in differentiating conflicts between the self and other (28, 42, 43). Given the importance of the rTPJ in P-taking (1, 28, 42, 44), we were interested in further describing the neurocognitive processes involved.

As previous research has implicated the TPJ as a major network hub (22, 23), we hypothesised that the region would co-ordinate shifts in functional connectivity to other brain regions, via phase synchrony (45, 46). Indeed, we found that the rTPJ increased its phase-coupling to the medial PFC and posterior PCC – two regions also involved more generally in understanding the mental states of others (7, 20, 41) (i.e. mentalizing). We also found increased phase synchrony between the rTPJ and SMC, SMA, and PPC (Fig. 3), regions previously implicated in coding for the body schema, i.e. cortical correlates of the internal representation of the body and its postures and actions (12, 13). We propose that this functional link, which has been reported previously (47, 48), underlies the simulated rotation of the embodied self into another’s orientation and perspective, which so far has primarily been described behaviourally (1, 8, 10). The rTPJ also showed decreased phase synchrony with visual regions (VOTC), primarily the ventral stream of the right visual cortex, during high-level P-taking. Altogether, these findings can be interpreted as an active shift from externally-driven processing (i.e. bottom-up sensory information) to internal representations (i.e. self, body-schema rotation) during high level P-taking. This switch from processing external events to updating internal states and information has been previously linked with TPJ function (23, 49).

Taken together, these findings suggest that rTPJ acts as a hub for high-level perspective taking by routing visual information to internal representations of the self, the body and its action and posture repertoire, via theta-band phase synchronisation (Figure 4). This allows humans to actively project their own sense of self into another’s viewpoint, resulting in rapid and accurate perspective-taking responses (1, 8, 10, 11). Interestingly, the other side of the coin seems to be that aberrant activity in TPJ contributes to involuntary shifts in perspective, as experienced in so-called “out-of–body” experiences (OBE) (25, 26). This emerging framework is consistent with a recent model arguing that the TPJ acts as a “nexus”, hub, or convergence zone between different cognitive domains including social cognition, attention and executive function (31). Our results suggest that the TPJ plays an important role during complex social-cognitive processes like P-taking, by co-ordinating the activity between multiple brain regions and functional sub-networks into a coherent whole (28, 31, 32, 42) (Fig. 4). We further propose that theta oscillations could be the crucial network code for this integration process (1, 11, 32).

### Top-down Executive Processes during high-level perspective-taking

Along with the rTPJ, two additional regions showed significantly increased theta power with increasing angular disparity during P-taking (Fig. 2A): the lateral PFC, primarily overlapping with the right inferior frontal gyrus; and the right ACC. This theta-band activity was found during a slightly later period than the rTPJ, from 0.2-0.5 post-stimulus onset, suggesting that the ACC and lateral PFC contribute later to the process of P-taking. Interestingly, we also found that these two regions displayed directed functional connectivity, as measured by Granger causality, to the rTPJ, mediated by low frequency brain rhythms (1-10Hz), indicative of top-down processing (50).

Whilst activity within these two regions is typically associated with cognitive control (51) and conflict monitoring (52), they have also been implicated in a number of theory of mind studies (34, 53–55). Activity within this context has been argued to reflect the detection (7,56) (ACC) and resolution (33, 55) (lateral PFC) of conflict between self and other perspectives (57). We therefore propose (Fig. 4) that the connectivity from rACC and rPFC to rTPJ, during later stages of P-taking, reflects domain-general “top-down” executive control processes (58) required for suppressing the self-perspective, in favour of taking the other’s perspective (33, 34, 59), and/or for controlling the conflict between the physical self and the transformed self (the “other”) (1, 42, 60), allowing both representations to co-exist in the brain, similar to the experience of an OBE, where the self is located in two places at once (26). Whilst this interpretation of our results is based on substantial empirical research (51, 57), to avoid reverse-inference, future work could vary executive demands during P-taking (61), in combination with brain stimulation (1) to establish the causal role of the lateral PFC and rACC. Nevertheless, the observation that our effects in ACC and PFC were primarily related to theta oscillations, further corroborates the notion of top-down control, since theta has previously been shown to reflect top-down cognitive control processes involved in conflict monitoring (52) and error-related responses (62). Our complimentary finding that the ACC and lateral PFC exerted top-down influence on the TPJ via low frequency rhythms (1-10Hz, peaking in theta) is clearly consistent with ACC and PFC theta as a mechanism for cognitive control (63).

## Conclusion

This study examined the cortical networks involved in high-level social alignment (perspective taking), co-ordinated by theta-band oscillations. Low-frequency phase coupling in the theta-band, has previously been shown to contribute to the co-ordination of long-range neuronal interactions (64, 65), through which distributed neural assemblies become integrated into a coherent network (46). Our finding that theta-band phase-coupling synchronises the right temporo-parietal junction (rTPJ) to brain regions involved in theory of mind and regions coding for body schema supports this view, and suggests that perspective taking, and potentially other social cognitive processes, involve the co-ordination of spatially and functionally disperse brain regions via theta-band phase synchrony (64), further supported by low-frequency top-down influences from executive control areas (32).

## Materials and Methods

Additional details are provided in SI Materials and Methods.

### Participants

Data were collected from 18 participants (4 male, 14 female, mean age = 27.55, SD = 5.86). All participants had normal or corrected to normal vision and no history of neurological or psychiatric illness. All experimental procedures complied with the Declaration of Helsinki and were approved by the Aston University, Department of Life & Health Sciences ethics committee. Written consent was obtained from all participants.

### Experimental Paradigm and Design

The paradigm was adopted from a behavioural study by Kessler & Rutherford (8). The stimuli were coloured photographs (resolution of 1024 × 768 pixels), showing an avatar seated at a round table shown from one of four possible angular disparities (see Fig. 1, left: 60°, 160° clockwise and anticlockwise). In each trial one of the grey spheres on the table turned red indicating this sphere as the target. From the avatar’s viewpoint, the target could be either visible or occluded (VO) by a centrally resented black screen; or to the left or to the right (LR) inducing P-tracking or P-taking, respectively. Stimuli were presented in 12 mini-blocks of 32 trials, alternating between LR and VO conditions. On each trial participants were asked to make a target location judgement according to the avatar’s perspective by pressing the instructed key on an MEG-compatible response pad: the left key for “left” or “visible” targets from the avatar’s viewpoint and the right key for “right” or “occluded” targets. Accuracy feedback was provided after each trial in the form of a short tone. As in Kessler & Rutherford (8), we collapsed across clockwise and anticlockwise disparities, and separately collapsed correct responses for left and right and visible and occluded, respectively. This resulted in four separate experimental conditions (for two examples see Fig. 1, left): left/right judgements where the avatar is 160° from own perspective (LR-160);left-/right judgements where the avatar is 60° from own perspective (LR-60); visible/occluded judgments where the avatar is 160° from own perspective (VO-160); visible/occluded judgments where the avatar is 60° from own perspective (VO-60).

### MEG and Structural MRI Acquisition

MEG data were acquired using a 306-channel Neuromag MEG scanner (Vectorview, Elekta, Finland) made up of 102 triplets of two orthogonal planar gradiometers and one magnetometer. All recordings were performed inside a magnetically shielded room at a sampling rate of 1000Hz. Two participants had excessive head movement (>5mm), and were excluded from subsequent analyses. A structural T1 brain scan was acquired for source reconstruction using a Siemens MAGNETOM Trio 3T scanner with a 32-channel head coil (TE=2.18ms, TR=2300ms, TI=1100ms, flip angle=9°, 192 or 208 slices depending on head size, voxel-size = 0.8x0.8x0.8cm). More details are provided in SI Materials and Methods.

### MEG Preprocessing

All MEG data were pre-processed using Maxfilter to supresses external sources of noise from outside the head (66), and loaded into the Fieldtrip toolbox (67) for standard data cleaning and pre-processing steps (see SI Materials and Methods). The pre-processed data were then separated into the four experimental conditions and downsampled to 250Hz to aid computation time.

### MEG Source-Level

Following MEG-MRI coregistration (see SI Materials and Methods) source localisation was conducted using Dynamical Imaging of Coherent Sources (38) (DICS) which applies a spatial filter to the MEG data at every voxel of a canonical 0.8 cm brain-grid, in order to maximise signal from that location whilst attenuating signals elsewhere. The spatial filter was calculated from the cross-spectral densities for a time–frequency tile centred on the effects found at sensor level (3-6Hz; 0–650ms; gradiometer channels only; see Fig. 1, top-right; Supplementary Figure S1). For all analyses, a common filter across baseline and active periods was used and a regularisation parameter of lambda 5% was applied. Cluster-based non-parametric permutation testing was used to correct for multiple comparisons across voxels (35), for the LR-hard>LR-easy and VO-hard>VO-easy contrasts. The resulting whole-brain statistical maps were spatially smoothed using a robust smoothing algorithm (68) as implemented in bspmview, and presented on a 3D cortical mesh using the Connectome Workbench software (36). Using the spatial filters computed during source analysis, we extracted trial-by-trial time-courses from three regions of interest, as shown in Fig. 2A, using the MNI co-ordinates with the highest t-value within each region (Table S3).

### Granger Causality Analysis

The directed functional connectivity between these three ROIs was estimated using spectrally-resolved non-parametric Granger causality (37) as implemented in the Fieldtrip toolbox (67). Intact and scrambled time-series were split into 0.325s epochs to enhance the accuracy of the results (0-0.65s post stimulus onset), followed by Fourier transformation (Hanning taper; 2Hz spectral smoothing), before being entered into a non-parametric spectral matrix factorisation procedure. Granger causality was then estimated between each ROI pair and each ROI-scrambled time-series. Statistical analysis was performed using cluster-based permutation testing (35). Granger causal influence between two regions (A & B) was deemed significant if values were i) significantly greater than scrambled data (p<.05) and ii) significantly greater in one direction than another (i.e. A-to-B versus B-to-A, p<.05).

### Theta-band Imaginary Coherence

To estimate patterns of whole-brain connectivity supporting high-level perspective taking, mediated by the right TPJ, we quantified theta-band phase synchrony during LR-160 trials compared with LR-60 trials. A complex-valued spectral estimate at 5±2Hz for each grid-point was estimated using an adaptive spatial filter [the ‘PCC’ method, as implemented in *ft_sourceanalysis* (67)]. Coherence was used to quantify the phase consistency between a seed region in TPJ (MNI co-ordinates [40-58 36]) and every other voxel of the canonical 0.8 cm brain-grid, using *ft_connectivityanalysis* (67). Coherence values are normalised to range from 0 (no phase synchrony) to 1 (completely phase synchronised). We opted to project the complex-valued coherency estimates onto the imaginary axis, as suggested by Nolte et al., (39). This removes estimates of instantaneous phase, thereby reducing the influence of spurious connectivity resulting from MEG field spread (39). Further details on the quantification of “imaginary coherence” can be found elsewhere (39, 67). Whole-brain coherence maps from LR-160 and LR-60 trials were baseline-corrected and compared using cluster-based permutation-testing as implemented in the Fieldtrip toolbox (67). The resulting whole-brain statistical maps were spatially smoothed using a robust smoothing algorithm (68) as implemented in bspmview, and presented on a 3D cortical mesh using the Connectome Workbench software (36).

### Analysis Code

MATLAB code for all analyses is openly available online at https://github.com/neurofractal/perspective_taking_oscillatory_networks (69).

## Acknowledgements

We wish to thank Gerard Gooding-Williams for MRI acquisition and the Wellcome and Dr. Hadwen Trusts for supporting MEG scanning costs. RS was supported by a Cotutelle Ph.D. studentship from Aston University and Macquarie University.

## Supporting Information (SI) – Oscillatory Networks Of High-Level Mental Alignment: A Perspective-Taking MEG Study

### SI Materials and Methods

#### MEG Data Acquisition

Five head position indicator (HPI) coils were applied for continuous head position tracking, and visualised post-acquisition using an in-house Matlab script. Two participants had excessive head movement (>5mm), and were excluded from subsequent analyses. For MEG-MRI coregistration purposes three fiducial points, the locations of the HPI coils and 300-500 points from the head surface were acquired using the integrated Polhemus Fastrak digitizer. Visual stimuli were presented on a projection screen located 86cm from participants, and auditory feedback through MEG-compatible headphones. Data acquisition was broken down into three sequential runs, each lasting 8-10 minutes.

#### MEG Data Preprocessing

All MEG data were pre-processed using Maxfilter (temporal signal space separation, .96 correlation), which supresses external sources of noise from outside the head (1). To compensate for head movement between runs, data from runs 2 and 3 were transformed to participant’s head position at the start of the first block using the *–trans* option of Maxfilter. For each participant, the entire recording was band-pass filtered between 0.5-250Hz (Butterworth filter) and band-stop filtered to remove residual 50Hz power-line contamination and its harmonics. Data were then epoched into segments of 2500ms (1000ms pre, 1500post stimulus onset) and each trial was demeaned and detrended. Trials containing artefacts (SQUID jumps, eye-blinks, head movement) were removed by visual inspection, resulting in an average of 90.1 trials per condition, per participant (additional descriptive statistics reported in Table S1). For sensor-level analyses, ICA was used to identify and reduce residual EOG and ECG artefacts. Four MEG channels containing large amounts of non-physiological noise were removed from all source-level analyses.

#### MEG-MRI Coregistration

MEG data were co-registered with participants’ T1 MRI structural scan by matching the digitised head shape data with surface data from the structural scan (2). Subsequently, the aligned MRI-MEG image was used to create (i) a forward model based on a single-shell description of the inner surface of the skull (3), using the segmentation function in SPM8 and (ii) spatial normalisation parameters to create individual volumetric grids. To facilitate group analysis, each individual volumetric grid was warped to a template based on the MNI brain (8mm resolution). Subsequently the inverse of the normalisation parameters was applied to the template grid, for subsequent source analysis.

#### Behavioural Statistical Analysis

Behavioural reaction times (RT) from the experimental paradigm were extracted from E-Prime^®^ data files and converted to .*csv* format. Data from two participants with MEG movement over 5mm was discarded. All trials containing incorrect answers or response times greater than 2 standard deviations from the median were excluded from subsequent analyses. For the four experimental conditions (LR-160; LR-60; VO-160; VO-60), median RT from each participant were entered into a one-way ANOVA using the JASP statistics package. The raw statistical output is reported in Table S2.

#### Sensor Level Analysis

Sensor-level time-frequency representations (TFRs) were calculated using a single Hanning taper between frequencies of 1-30Hz in steps of 1Hz. The entire 2500ms epoch was used, with a sliding window of 500ms, but the first 250ms and last 500ms of each trial were discarded to avoid edge artefacts. Due to different scales between the two MEG sensor-types, only data from the gradiometers were used, with TFR power averaged across each pair post-hoc. All analyses were computed on single trials and subsequently averaged, and therefore TFRs contain both phase-locked (evoked) and non phase-locked (induced) information. As hypothesised from previous research using a similar paradigm (4), TFR responses averaged across subjects showed prominent differences between conditions within the theta-band (3-7Hz).

For statistical testing, we compared theta-band (3-7Hz) power during trials in which the avatar was 160° versus 60° from the participant’s own perspective (clockwise or anticlockwise), in both left/right judgements (i.e. P-taking), and visible/occluded judgements (i.e. P-tracking). We corrected for multiple comparisons across time, frequency and space via cluster-based non-parametric permutation testing (5). Results showed a significant cluster of greater (3-6Hz) theta-band power at 0-650ms in the LR-160 versus LR-60 condition (highlighted in Fig S1, left), but not for VO-160 vs. Vo-60, (Fig. S1, right).

#### Source Level Theta-Band Power (3-6Hz) During P-tracking

To investigate changes in theta-band power accompanying P-tracking, we used the Dynamical Imaging of Coherent Sources (6) (DICS) approach described in Materials and Methods, for 0-650ms post-stimulus onset. Whole-brain theta power (3-6Hz) maps were averaged across VO-160 and VO-60 trials and compared to theta power maps from the pretrial baseline period, 0-650ms pre-stimulus onset, using non-parametric cluster-based statistics (5). Results show an increase in theta-power within ventral occipital and parietal regions (Fig. S2). This is consistent with the role of theta oscillations during event-related processing of visual information and decision making (7, 8).

Next, to investigate a potential angular disparity effect during P-tracking, baseline-corrected theta power maps were compared between VO-160 and VO-60 conditions, using cluster-based non-parametric statistics (5). As hypothesised, there were no significant differences between theta-band power in the VO-160 and VO-60 conditions (p>.2, Fig. S3).

#### SI Figures

**Figure S1:**
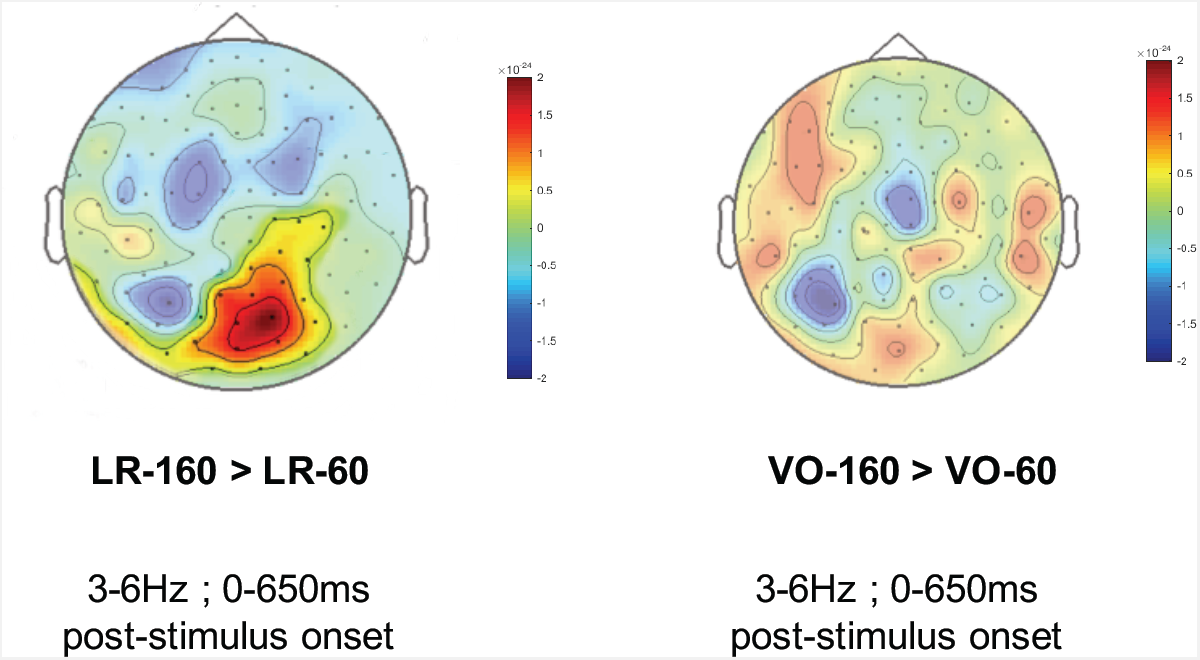
Sensor-level topoplots displaying oscillatory power between 3-6Hz, 0-650ms post-stimulus onset, for LR-160 > LR-60 and VO-160 > VO-60 contrasts. A significant cluster of greater (3-6Hz) theta-band power in the LR-160 versus LR-60 condition is highlighted (left), p=0.03 calculated using cluster-based non-parametric statistics (5).

**Figure S2:**
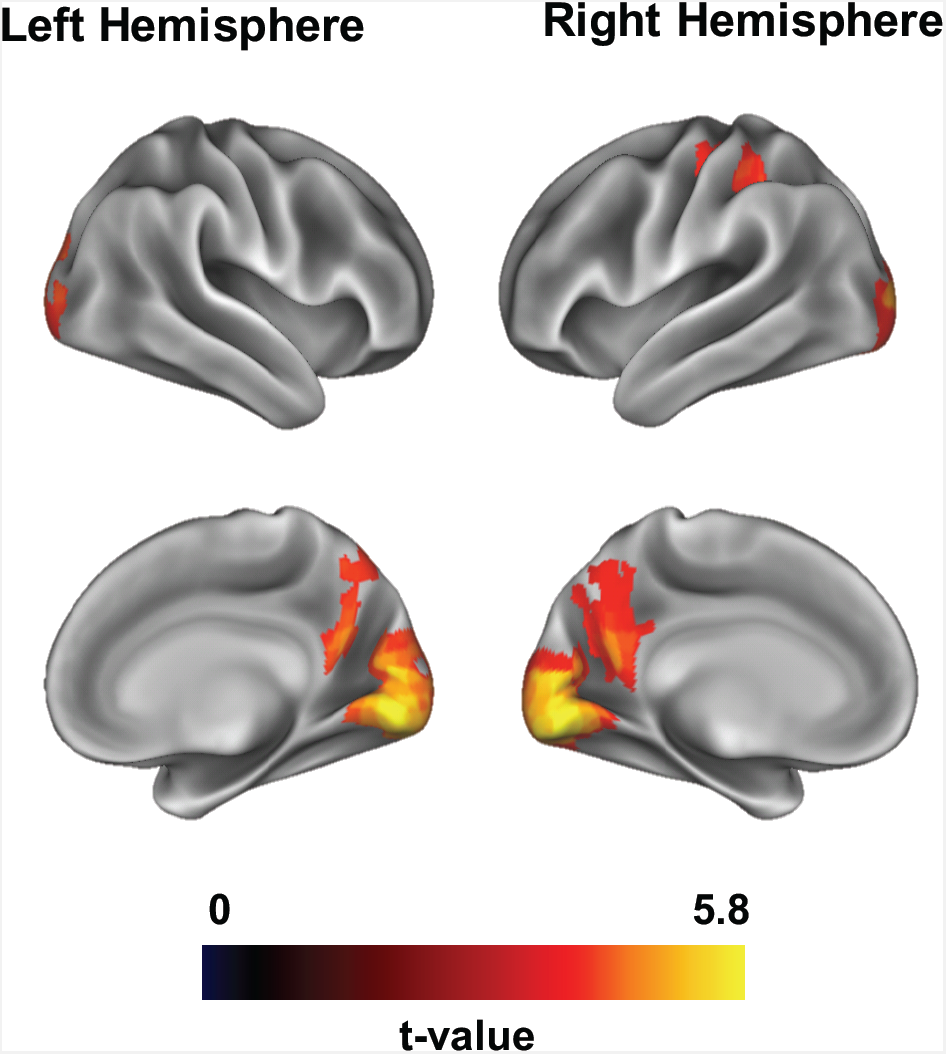
P-tracking results of the whole-brain DICS power analysis for visible/occluded > baseline contrast visualised using the Connectome Workbench software (9). Four significant clusters passing a p<.05 threshold were found within bilateral ventral occipital and right parietal cortices (5).

**Figure S3:**
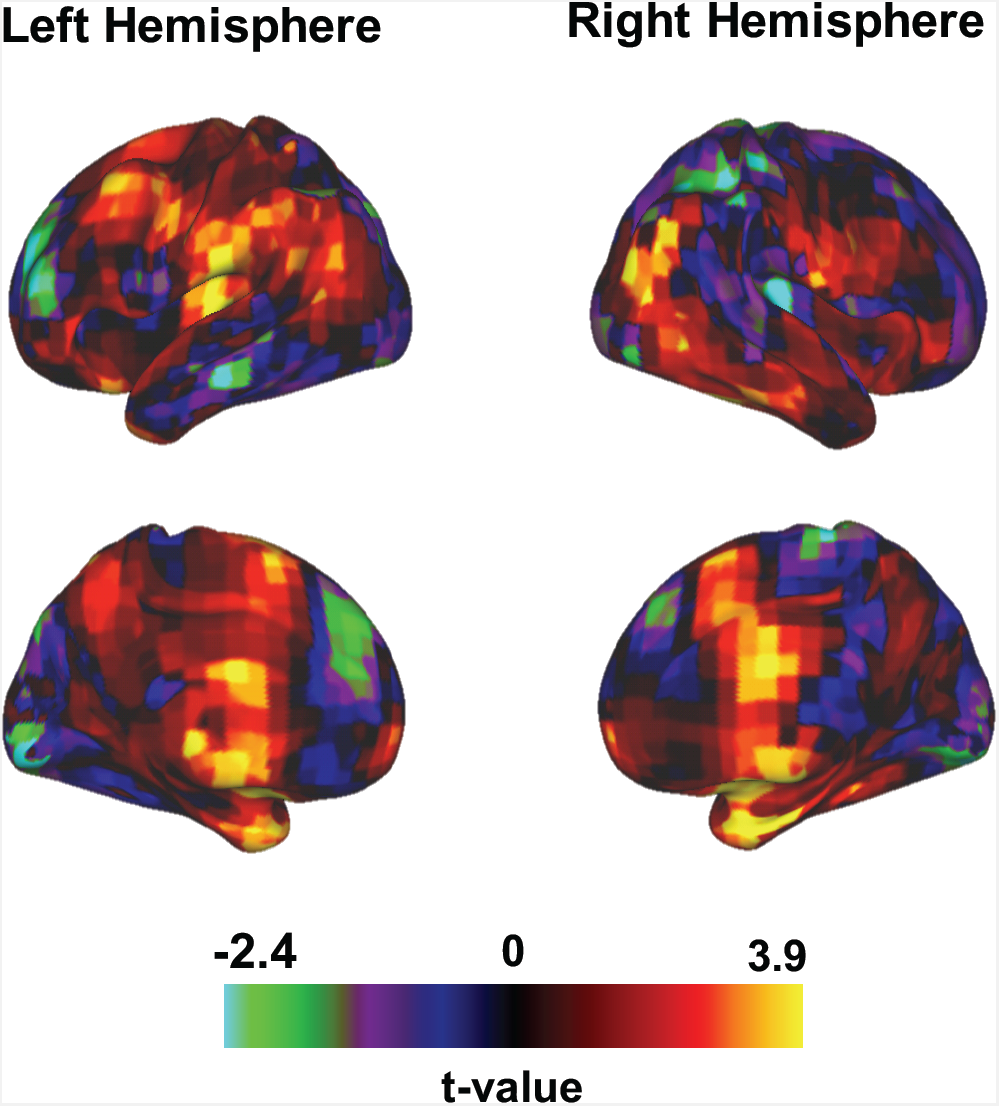
P-tracking results of the whole-brain DICS power analysis for VO-160 > VO-60 contrast visualised using the Connectome Workbench software (9). No significant clusters passing a p<.05 threshold were found (5).

#### SI Tables

**Table S1.** Descriptive statistics for the number of trials per condition, per participant, included in all MEG analyses.

| Conditon | Mean | Standard<br>Deviation | Max | Min |
| --- | --- | --- | --- | --- |
| LR_160 | 89.6 | 3.2 | 96 | 86 |
| LR_60 | 91.2 | 4.1 | 96 | 83 |
| VO_160 | 91.2 | 3.1 | 96 | 83 |
| VO_60 | 88.4 | 5.6 | 96 | 72 |

**ANOVA - RT\_time**
| Cases | Sum of Squares | df | Mean Square | F | p |
| --- | --- | --- | --- | --- | --- |
| Condition | 424154 | 3 | 141385 | 4.430 | 0.007 |
| Residual | 1.915e +6 | 60 | 31918 |  |  |

*Note.* Type III Sum of Squares

**Table S2.** JASP statistical output: a one-way ANOVA was conducted to investigate the main effect of condition on reaction time (RT). This was followed by post-hoc comparisons, using Tukey’s procedure to correct for pairwise comparisons.

|  |  | Mean Difference | SE | t | p<sub>tukey</sub> |
| --- | --- | --- | --- | --- | --- |
| LR-160 | LR-60 | 198.722 | 63.16 | 3.146 | 0.013 |
|  | VO-160 | 180.319 | 63.16 | 2.855 | 0.029 |
|  | VO-60 | 182.856 | 63.16 | 2.895 | 0.026 |
| LR-60 | VO-160 | -18.403 | 63.16 | -0.291 | 0.991 |
|  | VO-60 | -15.866 | 63.16 | -0.251 | 0.994 |
| VO-160 | VO-60 | 2.537 | 63.16 | 0.040 | 1.000 |

**Table S3.**
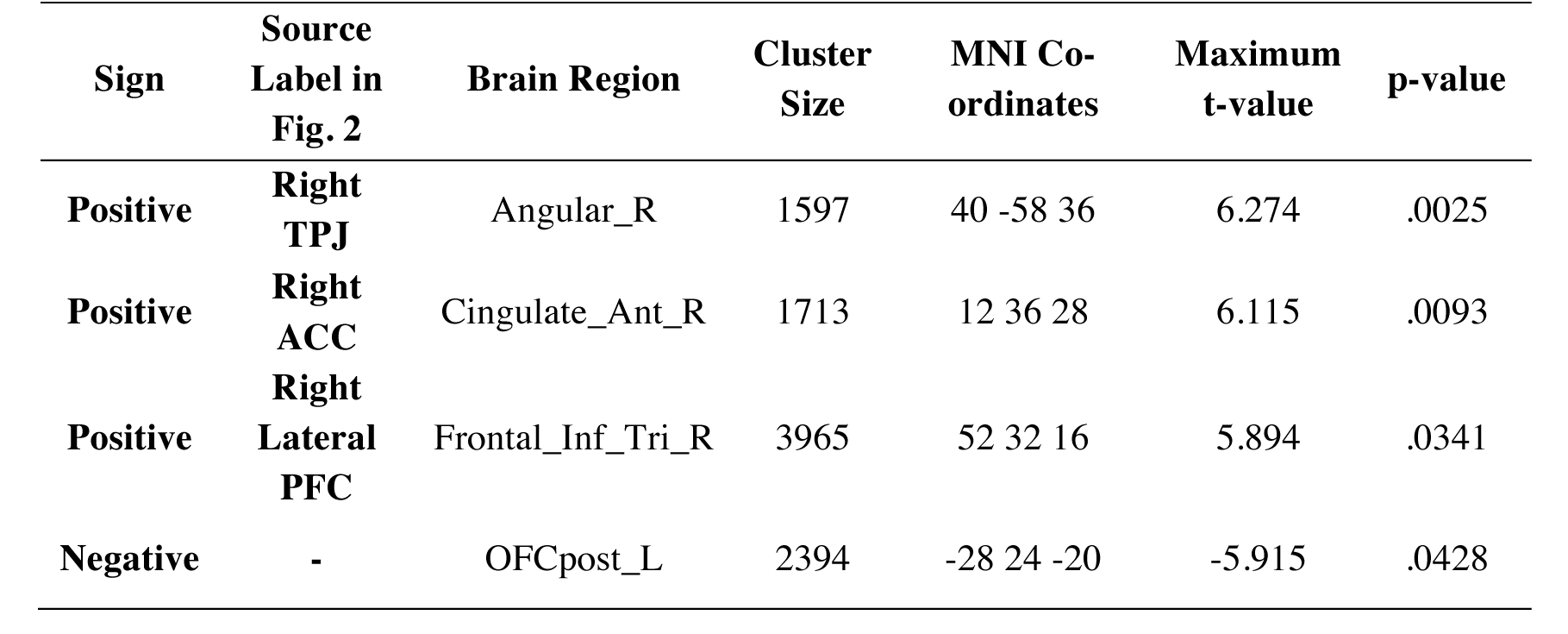
List of cortical sources for the LR-160>LR-60 theta-band power contrast shown in Fig. 2, Panel A in the main text. Table shows all local maxima separated by more than 20 mm. Regions were automatically labelled using the AAL2 atlas. Montreal Neurological Institute (MNI) coordinates correspond to the left-right anterior-posterior and inferior-superior dimensions respectively.

| Sign | Source Label in Fig. 2 | Brain Region | Cluster Size | MNI Co-ordinates | Maximum t-value | p-value |
| --- | --- | --- | --- | --- | --- | --- |
| Positive | Right TPJ | Angular_R | 1597 | 40 -58 36 | 6.274 | .0025 |
| Positive | Right ACC | Cingulate_Ant_R | 1713 | 12 36 28 | 6.115 | .0093 |
| Positive | Right Lateral PFC | Frontal_Inf_Tri_R | 3965 | 52 32 16 | 5.894 | .0341 |
| Negative | - | OFCpost_L | 2394 | -28 24 -20 | -5.915 | .0428 |

**Table S4.**
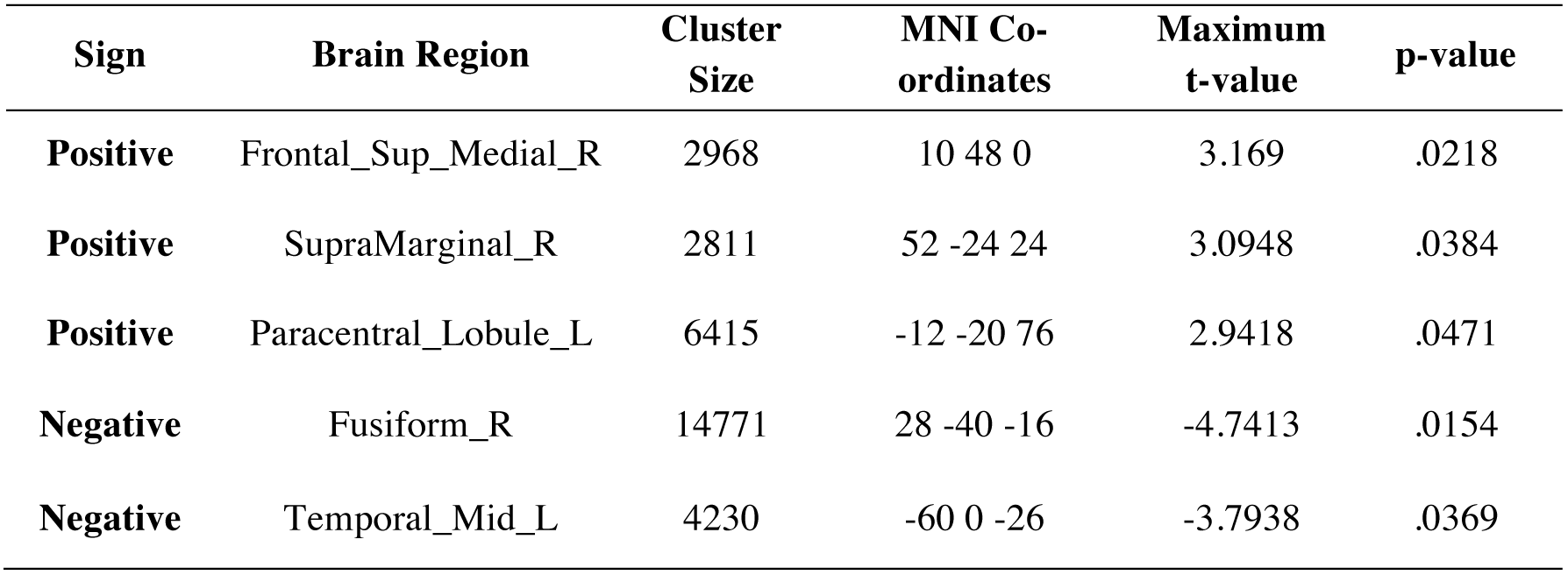
List of cortical sources for the LR-160>LR-60 theta-band imaginary coherence contrast shown in Fig. 3 in the main text. Table shows all local maxima separated by more than 20 mm. Regions were automatically labelled using the AAL2 atlas. Montreal Neurological Institute (MNI) coordinates correspond to the left-right anterior-posterior and inferior-superior dimensions respectively.

